# AI-Augmented Physics-Based Docking for Antibody-Antigen Complex Prediction

**DOI:** 10.1101/2024.11.06.622293

**Authors:** Francis Gaudreault, Traian Sulea, Christopher R. Corbeil

## Abstract

Predicting the structure of antibody-antigen complexes is a challenging task with significant implications for the design of better antibody therapeutics. However, the levels of success have remained dauntingly low, particularly when high standards for model quality are required, a necessity for efficient antibody design. Artificial intelligence (AI) has significantly impacted the landscape of structure prediction for antibodies, both alone and in complex with their antigens. We utilized AI-guided antibody modeling tools to generate ensembles displaying diversity in the complementarity-determining region (CDR) and integrated those into our previously published AlphaFold2-rescored docking pipeline, a strategy called AI-augmented physics-based docking. We highlight that the quality of the ensemble is crucial for docking performance, that including too many models can be detrimental and that prioritization of models is essential for achieving good performance. In this study, we also compare docking performance with AlphaFold, the new benchmark in the field. We distinguish between two types of success tailored to specific downstream applications: 1) criteria sufficient for epitope mapping, where gross quality is adequate and can complement experimental techniques, and 2) criteria for producing higher-quality models suitable for engineering purposes. Our results robustly demonstrate the advantages of AI-augmented docking over AlphaFold2, further accentuated when higher standards in quality are imposed. Docking performance is noticeably lower than the one of AlphaFold3 in both epitope mapping and antibody design. While we observe a strong dependence on CDR-H3 length for physics-based tools on their ability to successfully predict, this helps define an applicability range where physics-based docking can be competitive to AlphaFold3.

## Introduction

Predicting the structure of antibody-antigen complexes has tremendous value in biomedical research, with implications in understanding immune responses and designing therapeutics. While recent years have seen a growing reliance on sequence-based approaches for antibody research^1,2^, access to structural information remains crucial for applications requiring the spatial arrangements of protein atoms such as affinity maturation^3^ and developability improvement^4–6^. Accurate predictions of antibody-antigen complexes at atomic level remains a challenging task. Predicted structural models often come with uncertainties, emphasizing a persistent need for developing more accurate modeling tools. Recent advancements in deep learning, notably with the emergence of AlphaFold^7,8^, have renewed the optimism for improvements in protein structural prediction. Although AlphaFold has demonstrated an unprecedented performance in predicting generic protein-protein complexes, it has shown lower success rates against antibody-antigen complexes^9^, primarily due to the absence of co-evolutionary constraints.

Alternative methods exist to predict structures of complexes such as traditional physics-based molecular docking tools^10–12^. In the realm of molecular docking, a prevalent limitation arises from the lack of appropriate simulations of protein backbone flexibility, thus constraining the accuracy and the applicability of docking^13^. The significance of this limitation is underscored in the context of the complementarity-determining region (CDR) of antibodies, and particularly the hypervariable CDR-H3 loop, a region known for its conformational diversity and critical role in antigen recognition^14,15^. While some successful applications of protein docking have been reported for more rigid systems^16^ or when experimental information is leveraged^17^, it has generally shown limited success for more conformationally-diverse proteins, which are inherently more difficult to model, such as antibodies.

In response to these limitations, in a previous study we explored integrating traditional docking methods with AlphaFold^18^. Physics-based docking tools were used to generate a plethora of structural models, which were then evaluated by AlphaFold, similarly to other studies in related fields^19,20^. Reminiscent of rescoring techniques^21,22^, models were re-prioritized using the confidence scores of AlphaFold. The combination of physics-based methods with AlphaFold was shown to outperform AlphaFold-Multimer in antibody-antigen docking scenarios while being less demanding in resources, highlighting the benefits of template-based approaches over template-free modeling approaches. Moreover, we underlined the transformative capability of AlphaFold in remodeling the backbone during the rescoring process, which often improved the quality of the model. This feature not only addressed some of the limitations associated with backbone rigidity but also introduced a valuable compensatory mechanism mitigating the search-scoring trade-off in traditional rigid-docking methods. This advancement marked a significant stride towards overcoming the constraints of rigid docking protocols, and thus it holds promise for more realistic and biologically relevant predictions in the intricate atomic-level landscape of molecular interactions.

Recent years have witnessed a growth in the number of deep-learning-guided antibody modeling tools that can predict antibody structures in their free or unbound states^23–25^. While it is a matter of debate whether models of unbound antibodies are valid for antigen docking, there is increased evidence that bound conformations of antibodies are sampled in their unbound states^26–28^. The performance of antibody modeling tools in reproducing binding-competent states on untrained data has significantly improved over the years^23^. The more traditional antibody modeling tools relied on energy-based scoring functions^29,30^ that often failed to discriminate correct models from incorrect ones. One interesting feature of the newer AI-guided modeling tools is their ability to provide models with confidence estimates of the expected error to the ground truth. However, the trustworthiness of confidence estimates has not been clearly demonstrated for traditional physics-based docking models, which generally have been more sensitive to fluctuations in protein backbone conformation.

In a previous performance assessment of AI-enhanced physics-based docking tools, we relied on antibody-antigen systems in which both the bound and the unbound antibody structures were available from X-ray crystallographic studies. This greatly reduced the number of systems available for benchmarking and impacted the statistical significance of that study. Moreover, the full potential of AI-enhanced physics-based tools may not have been exploited by including a single, experimentally-determined, conformational state of the antibody CDR and thereby imposing rigid-backbone constraints during docking. Generating multiple structural models of a given antibody for antigen docking would better reflect many real-life applications which start from the antibody amino-acid sequence. To this end, the current study explored AI-generated ensembles of antibody models for use in antigen docking. We benchmarked performance against the latest cutting-edge tools available in this fast-evolving field^31^. As the antibody-antigen docking methods continue to improve, we emphasize the importance of success metrics in model evaluation and selection for subsequent antibody optimization.

## Methods

### Template-based modeling using docking

We collected antibody-antigen complexes from SAbDab^32^. Only entries that were released from January 1st, 2023 and onwards were retained to avoid any overlap with the data that was used to train the software used in this study. A resolution better or equal to 3.0 Å in the crystal structure was imposed. A total of 81 complexes met our criteria and were used as part of the benchmarking. The complete sequences were made available as part of the Supplementary Information.

The antibody models were produced by IgFold^24^, ABodyBuilder2^23^ and EquiFold^25^. The tools were run to obtain four antibody models per software. For ABodyBuilder2 and IgFold, one antibody model was predicted for each of the four independently-trained deep-learning models. For EquiFold, a single deep-learning model is available. The target of four antibody models was reached by re-running the tool multiple times producing a distinct conformation at every iteration. The antibody models were then energy-minimized with OpenMM^33^, PyRosetta^34^ and Amber^35^ for ABodyBuilder2, IgFold and EquiFold, respectively. The models were attributed a certainty value by standardizing the confidence values outputted by the software. The standardization was performed separately for ABodyBuilder2 and IgFold models across the whole panel of antibody systems. No certainty value was attributed to EquiFold-generated models from the lack of confidence values.

The antigen models were first obtained by dissociating the antigen from the antibody-antigen crystal structure. To ensure deviation from the bound state, perturbations in the spatial arrangements of side-chains were introduced using SCWRL4^36^. This step aimed to simulate a more realistic unbound state of the antigen.

The complexed states of the antibody-antigen were generated using protein-protein molecular docking software. ProPOSE^10^ version 1.0.3 and ZDOCK^11^ version 3.0.2 were run to generate the top-100 ranked models per software. The molecules were prepared as previously described^18^. The docking protocols were run with standard settings while constraining binding to the CDR region of antibodies. No epitope information was provided.

The antibody-antigen models produced in docking were rescored with AlphaFold version 2.3.1 using the model_1_ptm model with default parameters as previously described^18^. The docking-generated models were re-ranked according the AF2_ExpComposite_ in Equation 1.

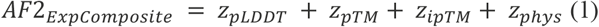

In comparison to the previous rescoring scheme^18^, the scheme in the current study was expanded to include the *Z*_ipTM_ and *Z*_phys_, which are the standardized ipTM and physics-based docking scores, respectively. As AlphaFold does not explicitly output the ipTM score when non-multimeric models are used, the value was manually calculated from the aligned confidence probabilities. All terms were weighted equally in the new scheme to avoid any potential overfit and allow for an increased transferability. The rescoring scripts are made available at the following repository: github.com/gaudreaultfnrc/AF2-Rescoring.

### Template-free modeling using AlphaFold

AlphaFold-Multimer^8^ was run through ColabFold version 1.5.2^37^ using the latest multimer_v3 models. For clarity, AlphaFold-Multimer is referred as AlphaFold2 in this paper. The AlphaFold2 calculations were run on Nvidia A100 Ampere, P100 Pascal and V100 Volta cards on the Digital Research Alliance of Canada clusters. AlphaFold version 3 (AlphaFold3)^31^ was run on the web server made accessible by its authors. While it would be ideal to run AlphaFold3 a thousand times with different seeds as recommended by their authors, it is simply not possible with the currently provided web server infrastructure. In an effort to minimize stochastic errors, AF3 was executed five times.

### Interface properties calculations

Starting from the crystal structures of the antibody-antigen complexes, surface areas (SA) of the residues were calculated. These surface areas were then compared to the fully exposed residues to determine the relative surface areas (rSA). The rSA was calculated for each residue in both the complexed (rSA_c_) and uncomplexed (rSA_u_) states, with the latter obtained by a rigid separation of the two molecules. The differences in rSA between these states were computed (ΔrSA). Residues were classified based on their rSA values as follows: interior (rSA_c_ < 25% and ΔrSA = 0%), surface (rSA_c_ > 25% and ΔrSA = 0%), support (rSA_c_ < 25% and ΔrSA > 0%), rim (rSA_c_ > 25% and ΔrSA > 0%), and core (rSA_u_ > 25%, rSA_c_ < 25%, and ΔrSA > 0%). The support, rim, and core regions were defined as the union of the residues in each respective class. Levy scores were calculated by summing the stickiness scores^38^ for all residues within these regions.

The global density (GD) serves as a proxy for determining the atom packing at the interface, specifically for atoms with a ΔrSA greater than 0%. To calculate GD, we first determine the volume of an ellipsoid based on the three moments of inertia from the subset of interface atoms. Then, the number of atoms is divided by the area of an ellipse defined by the two largest moments. The surface complementarity (SC) index is calculated by projecting normal vectors onto the tessellated surface of the interface. Normal vectors that are misaligned are penalized, and the resulting values are weighted by the tessellated area.

## Results and Discussion

Structural ensembles of free antibodies, i.e., without their antigens, were generated from sequence only for the 81 antibody-antigen systems in the test dataset. To generate conformational diversity in the CDR region of free antibodies, ensembles were modeled using a combination of three state-of-the-art AI-based antibody modeling tools (**Figure 1**). Four models were produced with each tool. For each system, we calculated the root-mean-square deviation (RMSD) values between models generated with each tool, called free-RMSD. The extent of CDR conformational diversity varied depending on the tool used. ABodyBuilder2 generated the most heterogeneous ensembles with free-RMSD values of 1.3 Å and 2.3 Å in the CDR and H3 regions, respectively. These values were accompanied by larger variations also in the framework region. In contrast, the ensembles produced by EquiFold were the most homogeneous, with free-RMSD values of 0.4 Å and 0.6 Å, respectively. The diversity among IgFold models was intermediate between ABodyBuilder2 and EquiFold.

**Figure 1.**
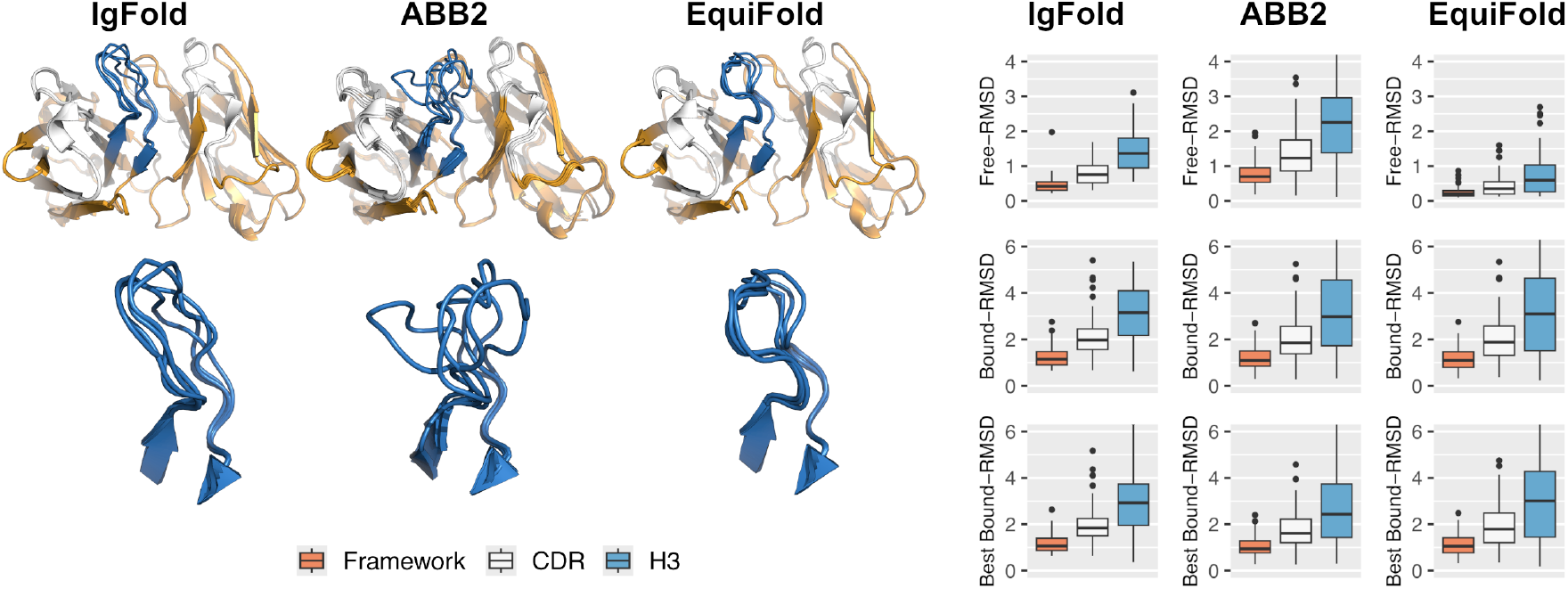
Structural diversity in the predicted antibody ensembles produced by IgFold, ABodyBuilder2 and EquiFold. The structural views depict representative deviations observed within the dataset for various antibody modeling tools. The free-RMSD describes the deviations among models per system and highlights the heterogeneous or homogeneous nature of the conformational ensembles generated by a given modeling tool. The bound-RMSD describes the deviations of free antibody models from the corresponding bound antibody structures. The RMSD values were calculated upon best-fit superposition of the framework region on the subset of backbone atoms.

To assess whether the ensembles could approximate the antibody state in the presence of its antigen, we calculated bound-RMSD values, which are relative to the bound states of antibodies as observed in crystal structures. When including all four models generated by IgFold, ABodyBuilder2, and EquiFold, the bound-RMSD values in the entire CDR were 2.0 Å, 1.9 Å, and 2.0 Å, and the bound-RMSD values in the H3 loop were 3.2 Å, 3.0 Å, and 3.3 Å, respectively. These relatively high values highlight the difficulty of modeling the bound states of antibodies from sequence alone. More noticeable differences between various tools emerged when evaluating the best models produced. ABodyBuilder2 generated models that were closest to the bound states, with overall bound-RMSD of 2.5 Å for the H3 loop compared to 2.9 Å for IgFold and 3.0 Å for EquiFold. The distribution for IgFold is noticeably tighter, signifying that there is less abundance of models closer to the bound state in comparison to EquiFold. These results suggest that ABodyBuilder2-generated ensembles might be better suited for physics-based docking studies due to their superior geometric complementarity to the antigen.

Using physics-based docking tools, we docked the antibody models against their respective antigens and rescored the top-100 predicted poses with AlphaFold2 as previously described^18^. Two different docking tools were used to explore potential benefits and synergistic effects of combining their predictions. Overall, 194400 antibody-antigen models were produced, equating to 32400 per free-antibody conformational ensemble per docking tool. We present the top-5 results based on two applications of docking: 1) for epitope mapping, where a general location of the binding site is sufficient, or 2) for antibody design, where more precise atomic resolution is required, especially at the antibody-antigen interface, to allow for further development such as affinity maturation. The two applications are associated with specific requirements for model quality. For antibody design, we imposed a requirement for at least a medium-quality model, defined as having at least 30% to 50% of native contacts, i.e., contacts observed in the antibody-antigen crystal. In contrast, for epitope mapping, we loosened the criteria and tolerated acceptable-quality models, defined as having at least 10% to 30% of native contacts.

First, we present docking results when no prior knowledge is available for model selection. In this instance, model selection must be performed naively by randomly choosing models. In this naive context, the maximum success rates were achieved using the EquiFold ensembles (**Figure 2**). We noticed minimal to no impact from using a single or multiple models for the free-antibody ensembles. Increasing further the ensemble size up to 10 models did not improve success rates (**Figure S1A**). The performance of AlphaFold2-rescored docking tools consistently outperformed AlphaFold2 alone, achieving success rates of up to 35% for epitope mapping. The performance drops slightly to 30% for antibody design. On the other hand, the performance of AlphaFold2 drops more importantly from 28% for epitope mapping to 13% for antibody design, while AlphaFold3 showed more stable performances of 47% and 46%, respectively. In both applications, the performance of AlphaFold3 exceeded the one of AI-augmented physics-based docking predictions, underscoring the substantial improvements in interface modeling from AlphaFold2 to AlphaFold3. When restricting the analyses to a single replicate, AlphaFold3 achieves equivalent numbers to a separate benchmark study^39^ with 42% and 35%. When using the IgFold and ABodyBuilder2 ensembles, performances were consistently lower and the impact on the number of models was more pronounced (**Figures S1B** and **S1C**). Interestingly, although ABodyBuilder2 produces the best antibody models on average (**Figure 1**), the inclusion of lower-quality models with higher bound-RMSD seems to introduce noise and reduce the performance. In line with these results, pooling the different ensembles into a single one impacted performance negatively (**Figure S1D**), implying a need for the prioritization of models.

**Figure 2.**
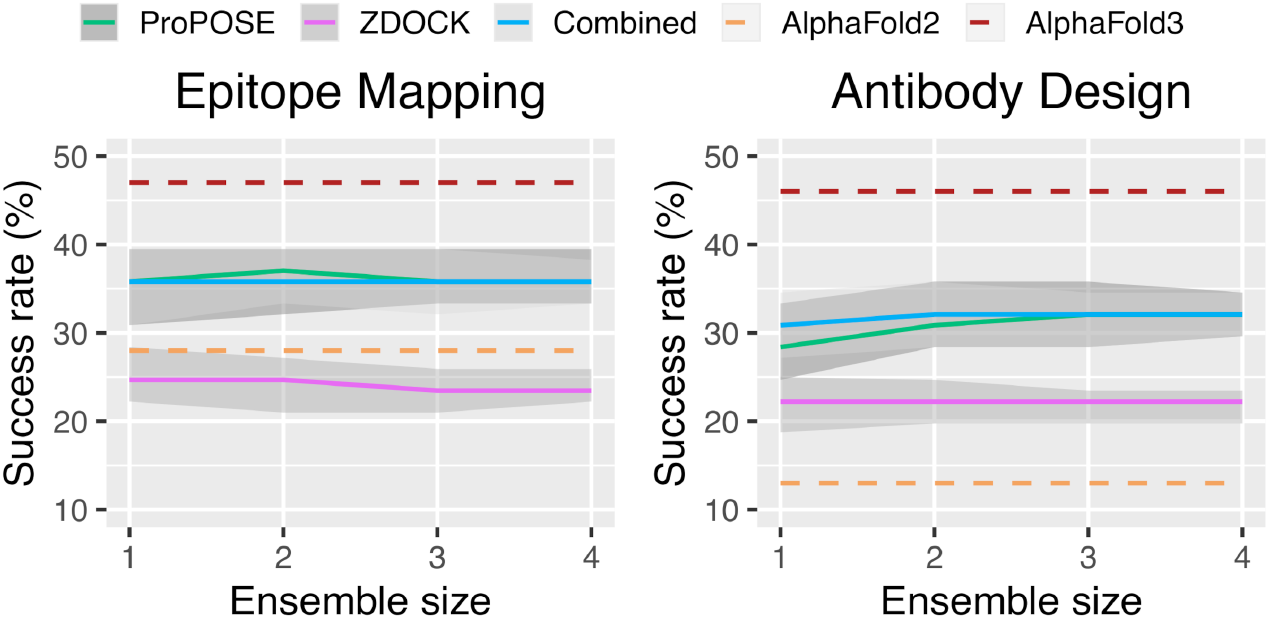
Success rates obtained from a naïve (i.e., random) selection of models. The rates were plotted as a function of the ensemble size to assess the impact of including an increasingly larger number of free-antibody conformational models. The performances of AI-augmented physics-based tools before (ProPOSE and ZDOCK) and after pooling their results (Combined) are compared to those of AlphaFold-Multimer (AlphaFold2) and AlphaFold3. The error bars were obtained from bootstrapping the antibody models with replicates for 200 iterations. The top-5 docking predictions using the EquiFold-generated models are shown.

Although the naive model selection approach is informative, it is not the most practical approach. Historically, physics-based functions were used to prioritize and select models based on energies, but these often failed to effectively discriminate between good and poor models. Instead, AI-guided modeling tools now provide us with confidence values for their predicted models, indicating how closely the models are projected to match the reference structures. We investigated the accuracy and reliability of the confidence values and explored how they could be leveraged for improved docking predictions. To facilitate the comparison between confidence scores among ensembles, we standardized the confidence scores on a scale of model certainty (**Figure 3**). We noticed a correlation between the length of the CDR-H3 loop with model certainty from the ABodyBuilder2 ensembles. While there is no guarantee that short CDR-H3 loops will be predicted with high confidence, the likelihood is relatively strong with a Pearson R^2^ of 0.58. Similar data were observed on the IgFold ensembles (**Figure S2**), but with weaker correlation (R^2^ = 0.36). A strong correlation was also found between CDR-H3 RMSD to the bound state, a proxy for model quality, with model certainty (R^2^ = 0.47) using the ABodyBuilder2 generated models. This suggests that model certainty could be used to prioritize models by filtering out poor-quality models. This may in turn reduce noise from the inclusion of too many models and thereby improve performance.

**Figure 3.**
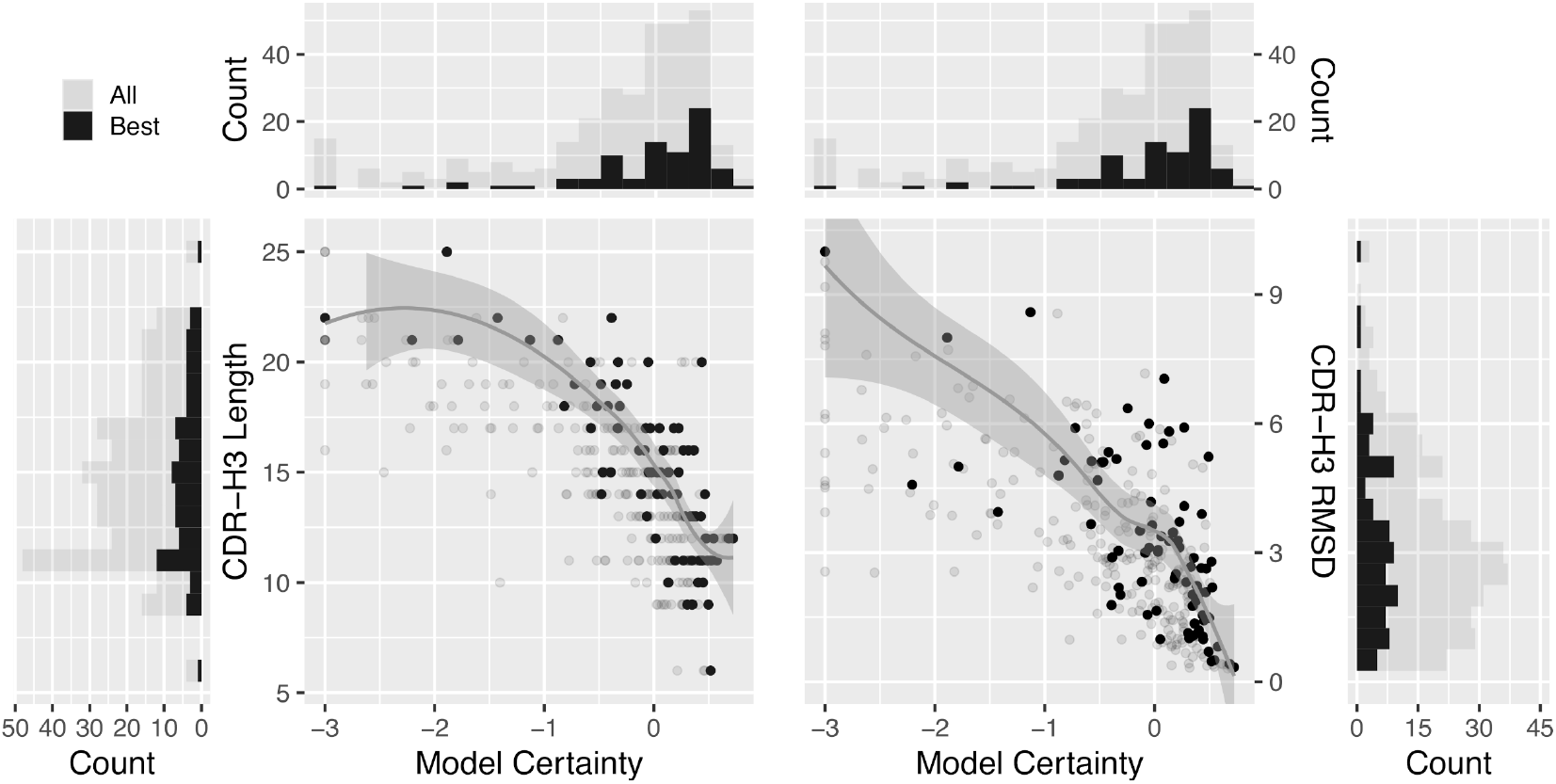
Standardized confidence scores plotted as model certainty against the length in CDR-H3 and RMS deviations to the CDR-H3 in the bound state, for the ABodyBuilder2 ensembles. The smoothed regression lines were built from the best subset of models, i.e. only considering the model with highest certainty for the antibody-antigen systems. The Pearson correlations (R^2^) for the best models are 0.58 and 0.47 for the CDR-H3 length and RMSD, respectively. Histograms were plotted to aid in visualizing the density of points in model certainty (top), CDR-H3 length (left) and CDR-H3 RMSD (right). Each bar corresponds to 0.2 certainty units, to 1 residue, or to 0.5 Å, respectively.

We tested this hypothesis and calculated the success rates while thresholding the input model quality to reject uncertain models predicted to be of lower quality. The performance in epitope mapping and antibody design are shown for the ABodyBuilder2 ensembles (**Figure 4**). With low stringency, success rates using the AI-augmented physics-based approach 31% in epitope mapping, slightly better than AlphaFold2 but still inferior to the naive selection from the EquiFold-generated models. We observed significant improvements in success rates as we increased the stringency of the threshold. While improvements in success were also noticeable using the IgFold ensembles, performance was generally lower (**Figure S3A**). Thresholding comes at a cost of reduced sample size: as the threshold becomes more stringent, more systems are excluded. In both epitope mapping and antibody design, the success rates reach those of AlphaFold3 when approximately 50% of systems are retained (N = 42), coinciding with an average CDR-H3 length of 12. At 25% systems remaining (N = 21), the performance of AlphaFold2-enhanced physics-based tools achieves 53% and 50% success in epitope mapping and antibody design, respectively. While it would be ideal for docking to be successful across the full range of CDR lengths, this is currently not feasible even with state-of-the-art antibody modeling tools. Nonetheless, our results highlight a specific applicability range for docking tools, providing valuable insights for future improvements.

**Figure 4.**
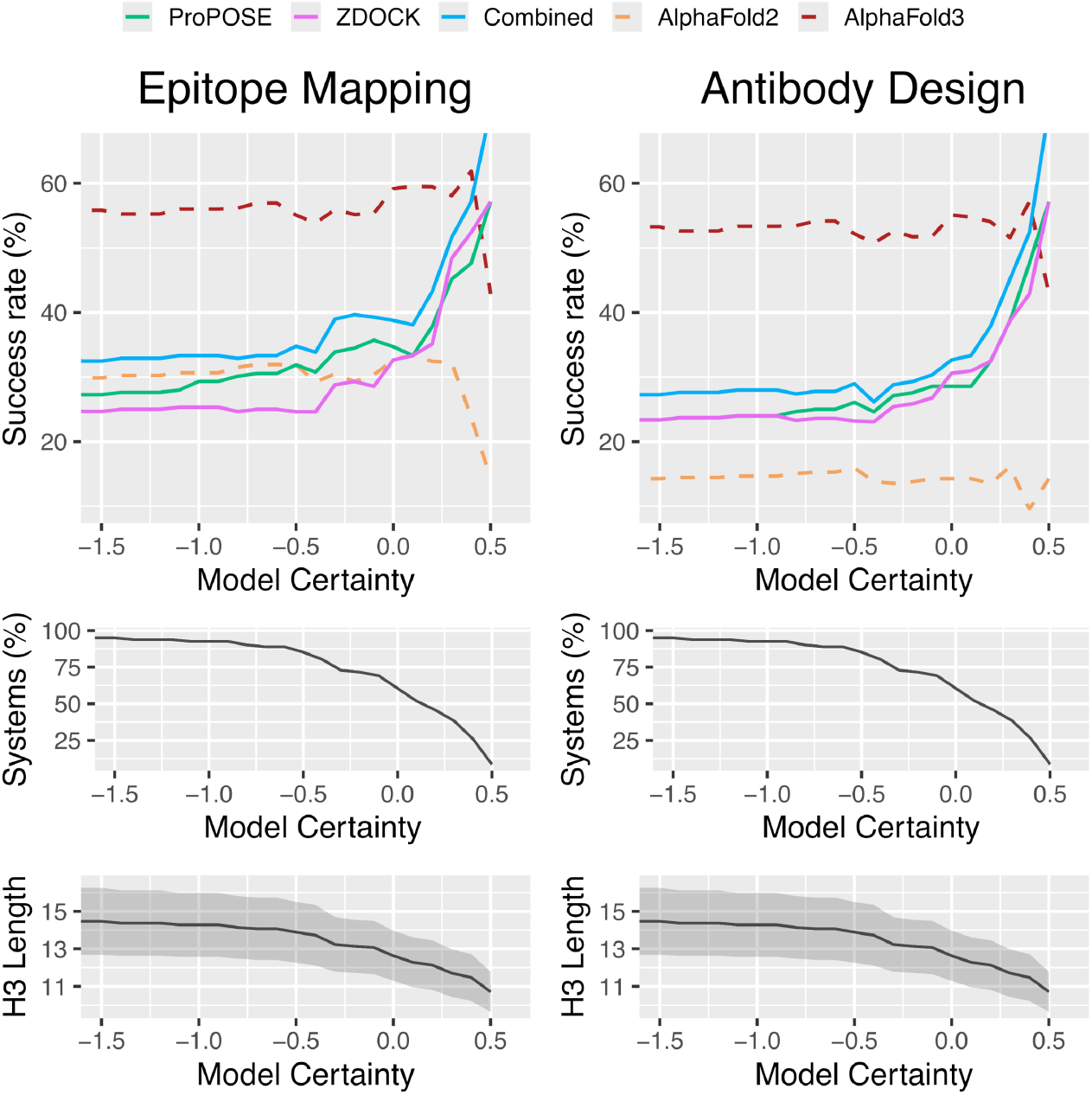
Success rates obtained from a confidence-guided selection of models. The rates were plotted as a function of the model certainty threshold below which antibody models are rejected. The performance of AI-augmented physics-based tools before (ProPOSE and ZDOCK) and after pooling their results (Combined) is compared to the ones of AlphaFold-Multimer (AlphaFold2) and AlphaFold3. For transparency, the number of systems remaining with their average length in CDR-H3 are reported. A minimum representation of 5% was imposed for data points to be plotted to minimize abruptness from the impact of low sample size. The top-5 docking predictions using the ABodyBuilder2-generated models are shown when using AbodyBuilder2-generated models.

While it may appear unfair to calculate success rates on different subsets of the dataset and compare those obtained by AlphaFold on the complete dataset, our approach mirrors how we would typically use AlphaFold, i.e. modeling from sequence alone without applying stringency based on an initial structural model quality. If we were to apply thresholding to AlphaFold in a similar manner, we would find that AlphaFold performance is not impacted on the different subsets. Provided the high correlation of model certainty with CDR-H3 length, this suggests that AlphaFold is less sensitive to CDR-H3 length than physics-based tools (**Figure S3B**) and that applying such a thresholding strategy would not enhance the performance for AlphaFold.

In an effort to find indicators for successful prediction, we characterized the antibody-antigen interfaces. We examined the whole interface as well as the support, core and rim regions that define the interface, according to previously described definitions^40^. These regions were analyzed in terms of size, through surface area calculations, and in terms of chemistry, through the stickiness score from Levy *et al*.^38^. Sticky interfaces are commonly observed in protein-protein interfaces due to their composition in amino acids, which is primarily characterized by the presence of hydrophobic groups. We complemented the analysis with the global density and surface complementarity index, properties that describe the packing of atoms at the interface, and the complementarity in molecular surface between the antibody and the antigen^41^. The properties were compared between successes and failures of the physics-based tools (**Figure 5**). It is clear that the amino acids that compose the binding interface is a strong determinant for success. Physics-based tools tend to fail more often for interfaces that are less sticky (p < 0.001), primarily from the core region of the interface (p < 0.01). Previous studies have raised the importance of surface area as a major contributor for success in rigid protein-protein docking, with smaller interfaces being more challenging to predict^10^. This trend becomes increasingly more evident as we increase stringency on model certainty, retaining only those models that approach their bound states and thereby mimicking closely the rigid docking experiment (**Figure S4**); the surface area properties all emerge as significant. The properties were also analyzed for successes and failures of AlphaFold3 (**Figure 5**). AlphaFold3 fails more often on systems with support regions that are less sticky (p < 0.01). The global density and surface complementarity index did not come up as significant determinants in the analyses.

**Figure 5.**
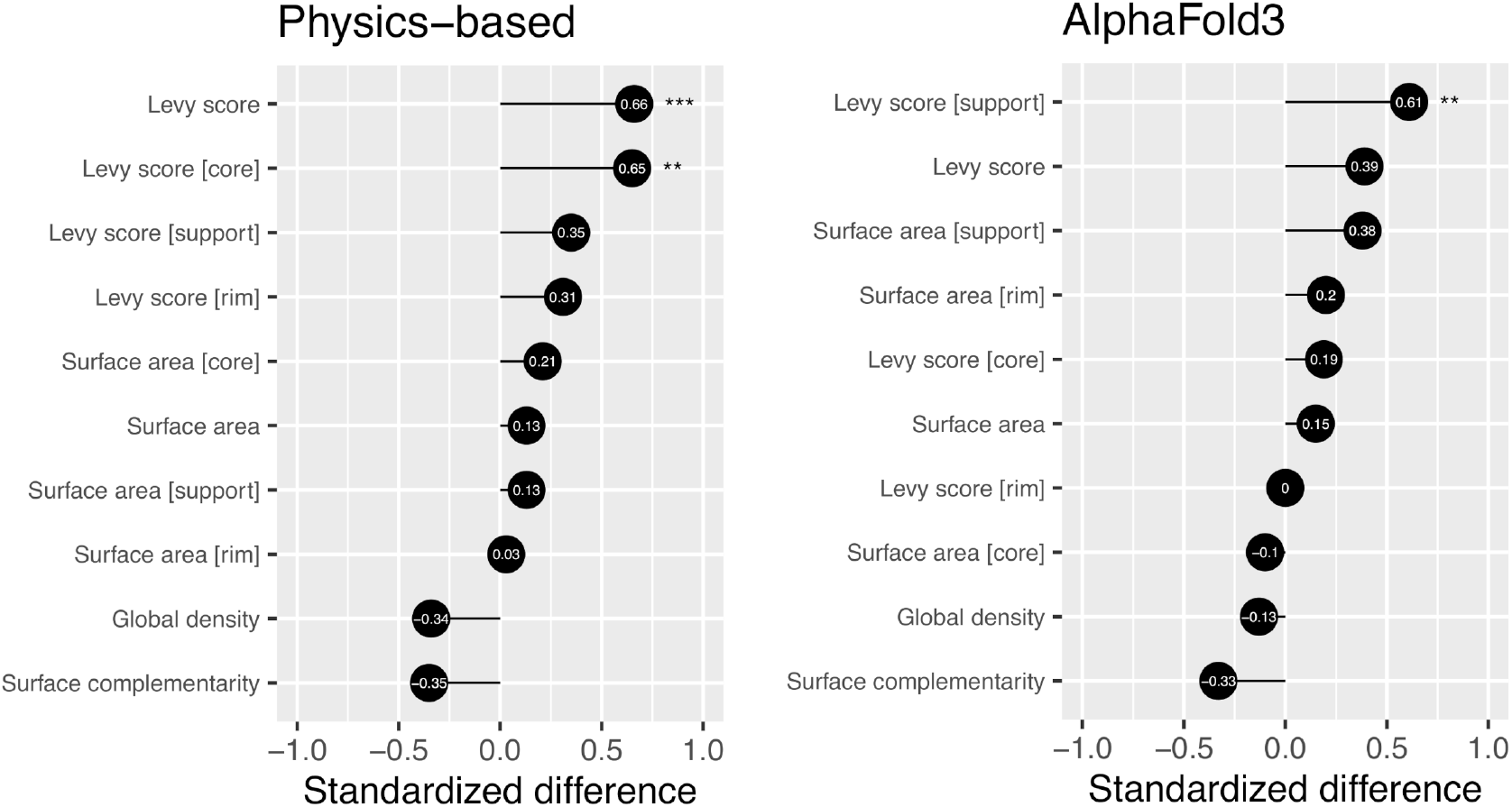
Standardized differences for a panel of properties characterizing antibody-antigen interfaces of crystal structures. The properties were calculated for models generated by physics-based tools and AlphaFold3 in the antibody design regime. The differences were calculated by comparing the mean of the subset of complexes that could be successfully predicted within the top-5 predictions to the mean of the subset of failures. Successes for physics-based tools are defined as the union of the successes across all tools used. P-values were calculated using t-tests from the underlying distributions of successes and failures. The significance of the p-values is indicated as follows: p < 0.05 (*); p < 0.01 (**) and p < 0.001 (***). Positive standardized difference values indicate higher success when the property is high and negative differences indicate higher success when the property is low.

## Acknowledgements

We thank Digital Research Alliance of Canada (formerly Compute Canada) for computing resource allocation for project number 4191.

## Author contributions

F.G., T.S. and C.C. contributed to the design of the study, the analysis and interpretation of the results. C.C. built and prepared the dataset. F.G. performed the bulk of the computational work. F.G. wrote the manuscript text and all authors contributed to the editing.

## Data availability

### Supplementary Information

accompanies this paper.

## Additional information

### Competing interests

The authors declare no competing interests.

## Notes

https://github.com/gaudreaultfnrc/AF2-Rescoring

### Competing Interest Statement

The authors have declared no competing interest.

