## Supplementary Information for "AI-Augmented Physics-Based Docking for Antibody-Antigen Complex Prediction"

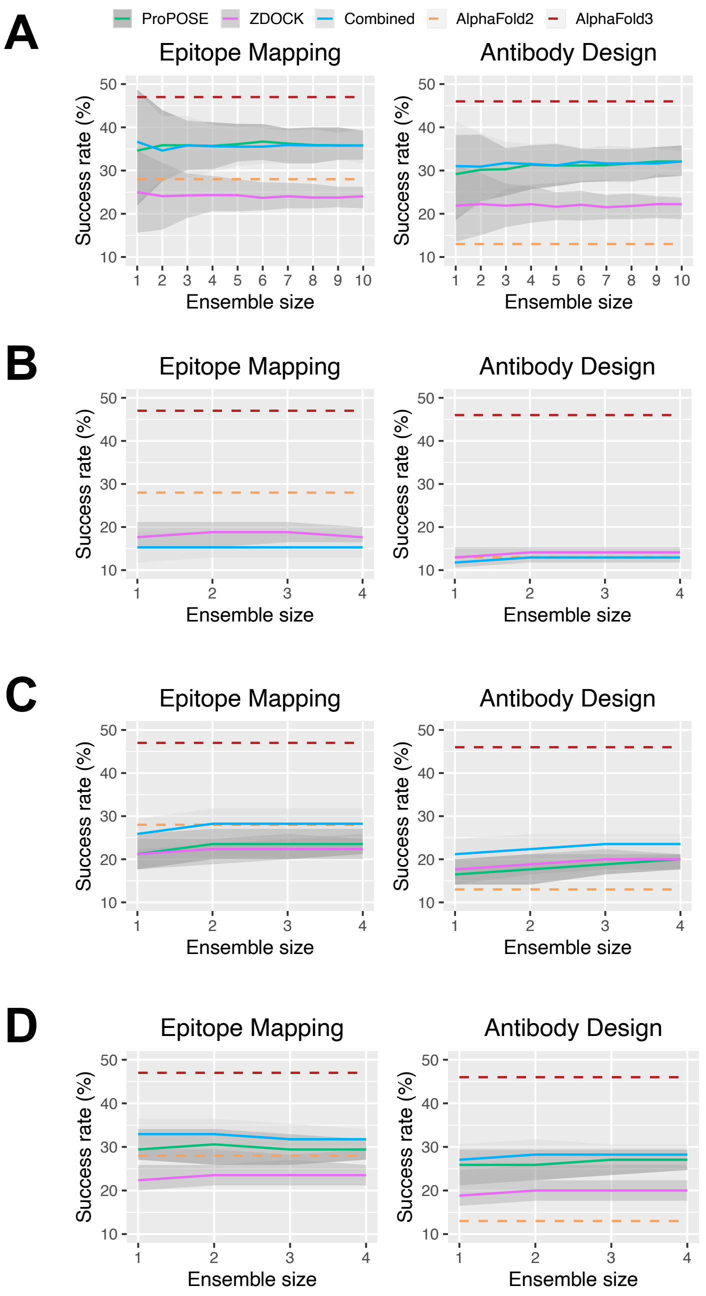


**Figure S1**. Success rates obtained from a naive selection of models. The rates were plotted as a function of the ensemble size to assess the impact of including an increasingly larger number of models. The performance of AI-augmented physics-based tools before (ProPOSE and ZDOCK) and after pooling their results (Combined) is compared to the ones of AlphaFold-Multimer (AlphaFold2) and AlphaFold3. The error bars were obtained from bootstrapping the antibody models with replicates for 200 iterations. The top-5 docking predictions are shown when using (**A**) the expanded EquiFold ensemble, (**B**) the IgFold-generated ensemble, (**C**) the ABodyBuilder2-generated ensemble and (**D**) the combination of the three ensembles.

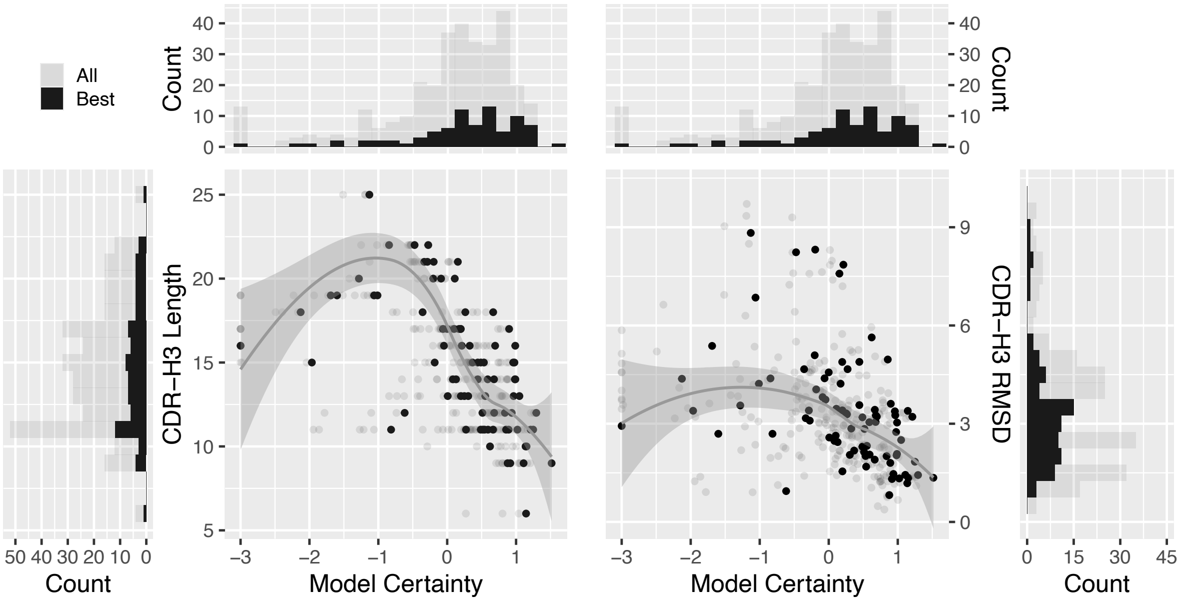


**Figure S2**. Standardized confidence scores plotted as model certainty against the length in CDR-H3 and RMS deviations to the CDR-H3 in the bound state, for the IgFold ensembles. The smoothed regression lines were built from the best subset of models, i.e. only considering the model with highest certainty for the antibody-antigen systems. The Pearson correlations ($R^2$) for the best models are 0.35 and 0.16 for the CDR-H3 length and RMSD, respectively. Histograms were plotted to aid in visualizing the density of points in model certainty (top), CDR-H3 length (left) and CDR-H3 RMSD (right). Each bar corresponds to one unit in 0.2 certainty, 1 residue and 0.5 Å.

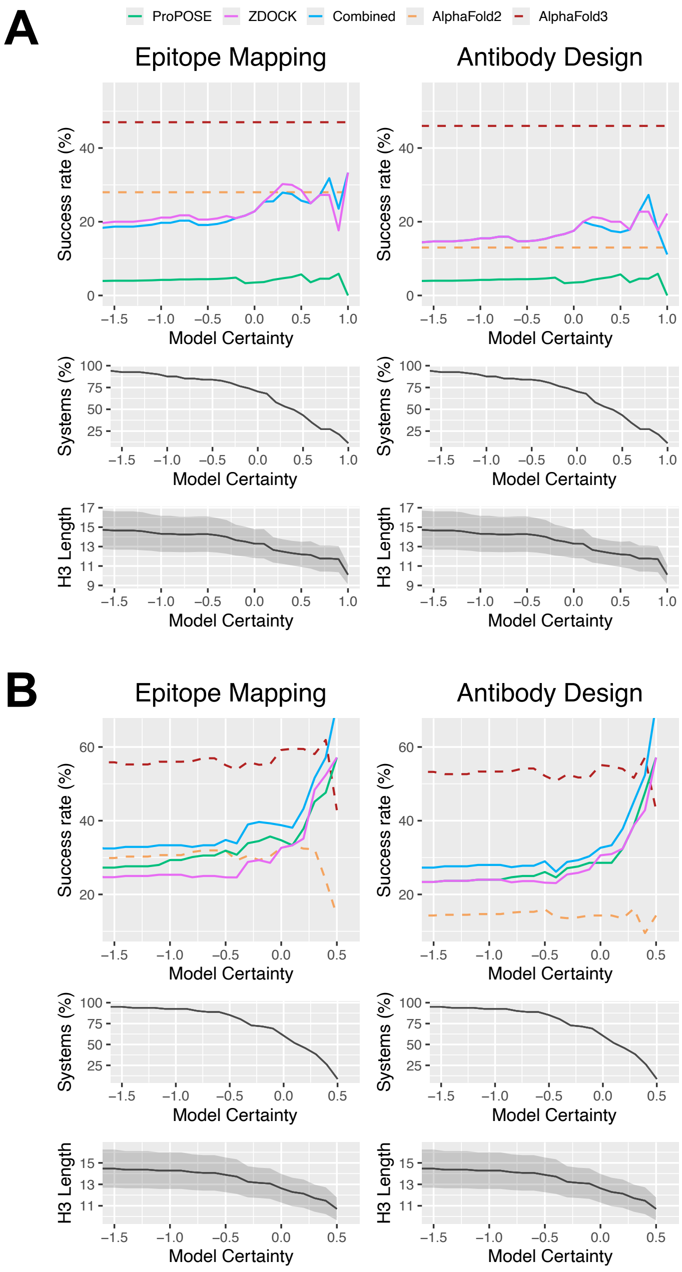


**Figure S3**. Success rates obtained from a confidence-guided selection of models. The rates were plotted as a function of the model certainty threshold below which antibody models are rejected. The performance of AI-augmented physics-based tools before (ProPOSE and ZDOCK) and after pooling their results (Combined) is compared to the ones of AlphaFold-Multimer (AlphaFold2) and AlphaFold3. For transparency, the number of systems remaining with their average length in CDR-H3 are reported. A minimum representation of 5% was imposed for data points to be plotted to minimize abruptness from the impact of low sample size. The top-5 docking predictions are shown when using (**A**) the IgFold-generated and (**B**) the ABodyBuilder2-generated models while thresholding to AlphaFold.

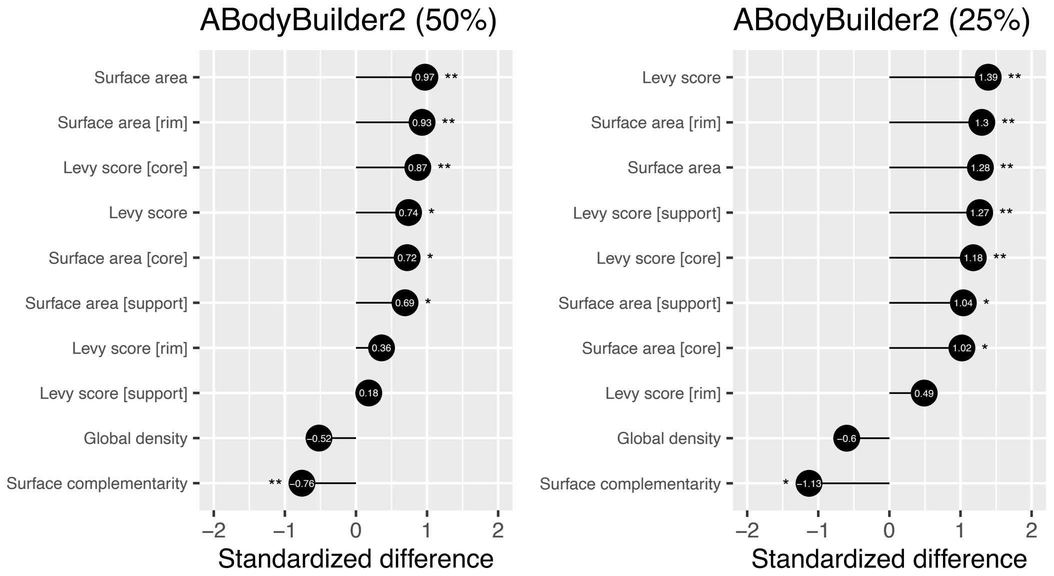


**Figure S4**. Standardized differences for a panel of properties characterizing antibody-antigen interfaces from crystal structures. The properties were calculated at 50% and 25% of remaining systems after excluding ABodyBuilder2-generated models with poor confidence. The differences were calculated by comparing the means between the subset of complexes that could be successfully predicted within the top 5 predictions to the subset of failures. Successes are defined as union of the successes across all physics-based tools used. P-values were calculated using t-tests from the underlying distributions of successes and failures. The significance of the p-values is indicated as follows: $p < 0.05$ (*); $p < 0.01$ (**) and $p < 0.001$ (***). Positive standardized difference values indicate higher success when that property is high and negative differences indicate higher success when the property is low.
